# Odor sensory tests vs. *In-silico* prediction for the high-definition quantification of olfaction dynamics

**DOI:** 10.1101/2024.10.12.617741

**Authors:** Islam M. S. Abouelhamd, Kazuki Kuga, Kazuko Saito, Megumi Takai, Takahiro Kikuchi, Kazuhide Ito

**Affiliations:** Interdisciplinary Graduate School of Engineering Sciences, Kyushu University, Kasuga-koen, Kasuga-shi, Fukuoka 816-8580, Japan; Faculty of Engineering Sciences, Kyushu University, Kasuga-koen, Kasuga, Fukuoka 816-8580, Japan; R&D Division, Meiji Co., Ltd, Tokyo, Japan

**Author notes:** Department of Architectural Engineering, Faculty of Engineering, Assuit University, Egypt (Islam M. S. Abouelhamd).

## Abstract

The intricate dynamics of volatile organic compounds (VOCs) in the human respiratory system remain poorly understood. In the present study, we integrate odor sensory tests (OSTs) coupled with computational fluid dynamics and a physiologically based pharmacokinetic (CFD-PBPK) model to elucidate various aspects of the olfaction process. Safe yogurt-derived substances were incorporated in OSTs to prevent harmful exposure. Acetaldehyde was identified as a key active component in determining odor intensity, prompting further analysis of acetone and other four constituents of yogurt. Logarithmic correlations were established between the perceived odor intensity from the OSTs and both time-averaged absorption flux and equilibrium concentration within the olfactory mucus layer. These parameters were numerically captured, enabling the logarithmic approximation of odor intensity for different breathing profiles and the development of reliable prediction models for odor sensation based on quantifiable physiological parameters. The CFD-PBPK model captured detailed spatial and temporal variations of these parameters, which offers potential for future integrated/in-silico applications. Minor peaks of odor concentration were observed in the posterior olfactory regions during exhalation, revealing a retro-nasal phenomenon. Location-specific analysis revealed the nostrils and olfactory regions as the most accurate indicators of perceived odor intensity, proving the limitations of rough sensory assessments in the indoor/breathing zone scales. Acetone exhibited distinct absorption and desorption trends during the transitional phase between inhalation and exhalation, owing to the physical properties (diffusion and partition coefficients) that strongly characterize the olfaction dynamics.

**Author Summary:** We developed an integrated method using odor sensory tests (OSTs) coupled with computational fluid dynamics and physiologically based pharmacokinetic models (CFD-PBPK) to assess the temporal and spatial transport of yogurt odorants to the human olfactory region. Multiple phenomena were observed, including ortho-nasal and retro-nasal olfaction, temporal changes in the perceived odor intensity associated with breathing/sniffing profiles, absorption and desorption curves of acetone in the mucus epithelium, and regional-based olfaction distribution. The perceived odor intensity from OSTs can be predicted logarithmically in correlation with both the time-averaged absorption flux and the equilibrium concentration in the olfactory mucus layer, offering a reliable *in-silico* prediction model for odor sensation based on numerically quantifiable parameters. This model offers potential implications for multiple computational, biomedical, and industrial applications, such as the electric noses, smart odor sensors, food assessments, and fragrance development, particularly for long-term exposure in the industries that emit odorous compounds. It can open the door for more accurate predictions of the complex micro-fluid dynamics in the microbial ciliated tissues in the olfactory receptors.

## Introduction

Human olfaction is fundamental in various biomedical and bioengineering applications. It plays an important role in determining communication, behavior, and food preferences [1, 2]. Although there are three key brain processes involved in odor identification, namely memorization of intensity, quality, and acceptability [3], the transport dynamics of volatile organic compounds (VOCs) to the respiratory tract remain poorly understood. Ethical concerns regarding human subject experiments, the complexity of *in-vivo* odorant quantification, and individual disparities further hinder research [4]. The distinguishing sensitivity of the olfactory receptors (ORs) in the mucus layer plays a crucial role in responding to VOCs through the interplay of olfaction and inhaled airflow biomechanics. Nevertheless, owing to indirect access to the olfactory recess, only 10% of inhaled air during normal breathing reaches the epithelium [5, 6], making it challenging to capture the target fluids and measuring reliable concentration profiles.

Following the proposed linear relation between the perceived air quality and the indoor acetone concentration [7], various contributions emerged to enhance odor sensation and promote healthy building principles [8–11]. While traditional research about odor intensity quantification relied on trained panelists [12], inconsistencies in these assessments have been widely debated [11, 13]. Recent studies suggest a logarithmic correlation between odor intensity and concentration [14]. However, the proposed functions depend on the scale of the indoor environment without considering the human airway domain. Thus, a deep understanding of fluid and mass transfer is required to unravel the underlying mechanism of olfaction. In this study, we used the odor sensory test (OSTs) associated with the human perception of odor intensity and acceptability, coupled with a computational fluid dynamics/physiologically based Pharmacokinetic Model (CFD-PBPK), to investigate multiple phenomena associated with the olfaction process. The VOCs from daily yogurt were adapted during the OSTs to ensure 100% safety for human subjects.

Convective diffusion transport to the mucus layer has numerous implications, including, but not limited to, diffusivity, solubility, partition coefficient, multiphase interactions, chemical reactions, breathing airflow characteristics, and anatomical features [15–23]. Earlier literature reveals that unsteady solutions are crucial for the accurate prediction of the uptake and deposition of chemical species over a wide solubility range [24,25]. High-solubility odorants are primarily absorbed in the anterior olfactory recess, whereas odorants with lower solubility exhibit more uniform absorption throughout the passage [26], aligning with gene expression zones [27]. Anatomical features influence olfaction [6]. Li et al. [21] reported individual disparities in olfactory sensitivity induced by the nasal cavity anatomy. Zhao et al. [28] reported a strong correlation between odorant detection thresholds (ODTs) and the minimal cross-sectional area of the olfactory region, accounting for 65% of the total ODTs in patients with olfactory loss due to chronic rhinosinusitis. Breathing/sniffing profiles can strongly characterize the airflow patterns in the olfactory recess [17, 18, 29]. Sniffing cycles can easily transport 2.5 times more air to the olfactory recess than quiet breathing, enhancing odorant uptake.

## Results and Discussion

We obtained a wide range of correlated parameters like perceived odor intensity (*I_o_)*, acceptability, equilibrium concentration (*C_ave_*), and absorption flux (*A_ave_*) (Figs 1A–1G), using two distinct methods to evaluate the olfaction process. These variables maximized the correlation analysis, enabling accurate prediction of odor intensity based on related quantifiable parameters. In the OST track, perceived odor intensity scores over five breathing and sniffing cycles reflected the sensitivity of the trained panelists to odor perception. The scores showed limited average intensity between 4.0 and 6.0 (out of a 10-point scale) during the peaks of the inhalation phases (Fig 1E). The cyclic intensity aligns with the cyclic nature of the respiration profile. Intensity peaks occurred in the second cycle during normal breathing, while sniffing cycles slightly surpassed the third cycle. However, due to the limitations of OSTs, the quantification of the odor concentration was not highly precise.

**Fig 1.**
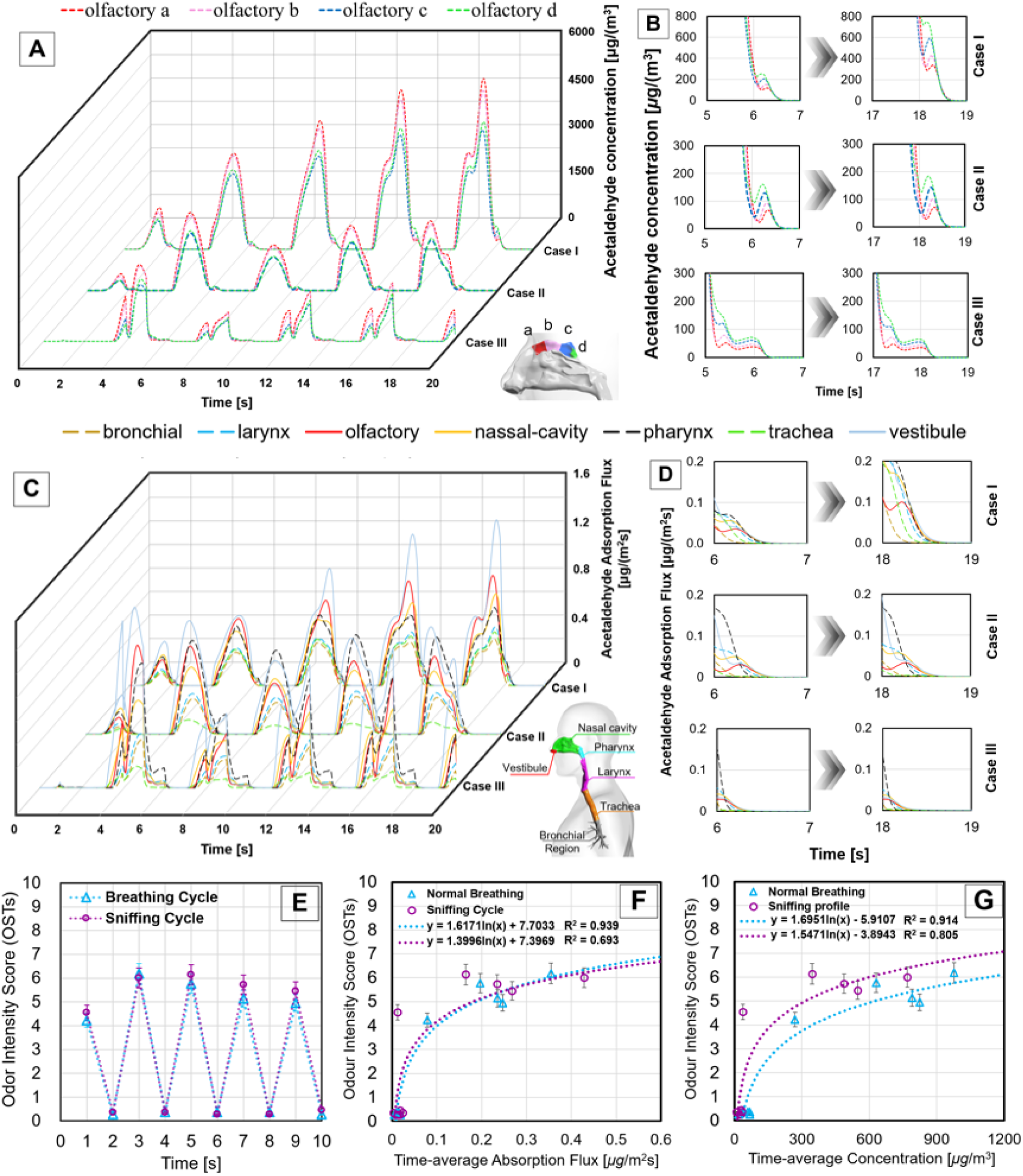
Perceived odor intensity and numerically quantified concentration/absorption flux using odor sensory tests (OSTs) and computational fluid dynamics and a physiologically based pharmacokinetic (CFD-PBPK) model, respectively. Panel (A) presents the temporal change of acetaldehyde concentration in the mucus layer of the olfactory region in three examined breathing profiles, while panel (B) clarifies the detailed trend of the retro-nasal olfaction phenomena during the exhalation periods, as predicted by the CFD-PBPK track. Similarly, panels (C) and (D) present the trend of absorption flux of acetaldehyde in the mucus layer in different airway segments highlighting the second peak observed in the olfactory region (D). Panel (E) shows the perceived odor intensity by the panelists for 20-second duration for five subsequent breathing and sniffing cycles. Panel (F) and (G) show the correlation analysis between the perceived odor intensity from the panelists during the OSTs and the time-averaged absorption flux and concentration which predicted from the numerical track, respectively.

The CFD-PBPK track can enhance our understanding of olfaction dynamics by detecting the absorption and concentration trends of individual chemicals in different temporal and spatial locations. Fig 1A illustrates the fluctuating acetaldehyde concentration (the primary odorant in yogurt) within four distinct segments of the olfactory recess (labeled as olfactory a, b, c, and d, from the anterior to the posterior regions). During inhalation, regardless of the breathing profile, anterior olfactory regions (olfactory a and b) received higher odorant concentrations due to direct interaction with the nostrils and nasal passages (ortho-nasal olfaction). However, during the transition from exhalation to inhalation, minor peaks were observed in the posterior olfactory segments (c and d); this can be attributed to the slight retro-nasal olfaction. The trapped odorants in the lower airway segments were released backward into the posterior olfactory recess, influenced by the exhalation airflow. This phenomenon, undetected by the OSTs, resulted in near-zero perceived intensity. Breathing profiles and cycle order influenced the different strength levels of this phenomenon, which gradually increased over multiple cycles, especially during normal breathing (Fig 1B). Regarding breathing profile, a high breathing flow rate (Case I) indicated a gradual increase of the acetaldehyde concentration in subsequent cycles, reaching a maximum of approximately 6,000 μg/m^3^ in the fifth cycle. Case II reported a lower overall concentration induced by a lower inhalation flow. Initially, low concentrations were detected in the olfactory region, followed by high concentrations in the second cycle. A subsequent drop in the third cycle was followed by a slight gradual increase. Case III, characterized by a powerful sniffing flow rate over a short period (one second), exhibited sharper peaks and quicker drops, suggesting rapid odorant uptake followed by swift clearance during exhalation. Thus, controlling the breathing profile is a fundamental factor for accurately predicting fluid and mass transfer. Check S1 – S5 Figs and S1 Table for detailed trends of concentration and absorption flux for different VOCs of the yogurt, which analyzed by the OSTs and CFD-PBPK methods.

The delineated absorption flux of acetaldehyde, as shown in Fig 1C, can provide additional insights into regional absorption trends. Each segment exhibits a distinct pattern, emphasizing the physiological roles of odorant uptake and clearance. Owing to proximity and direct exposure to the inhaled airflow, the mucus layers in the vestibule, olfactory, nasal cavity, and pharynx segments showed the highest periodic curves of the absorption flux, while the lower regions reported relatively lower values. Despite the location and intricate structure of the olfactory recess, a distinct trend in absorption flux emerged, characterized by multiple minor peaks during exhalation (Fig 1D). Another significant observation is the influence of airflow rate on the absorption flux. The higher trends in Case I compared to those in Cases II and III indicate that increased airflow leads to greater acetaldehyde uptake in the time-averaged value. However, the earlier absorption trends were associated with the sniffing cycle (Case III).

The correlation analysis between the outputs of OSTs and CFD-PBPK tracks, presented in Figs 1E, 1F and 1G, can approximate the odor intensity as a function of the numerically quantified time-averaged absorption flux [µg/m²·s] or concentration [µg/m³] of the chemicals in the mucus layer of the olfactory wall. For each perceived odor intensity (*I_o_*) by the panelists in each breathing cycle, representative values of the time-averaged (during the timeframe of each cycle), absorption flux (*A_ave_*), and equilibrium concentration (*C_ave_*) of acetaldehyde were determined from the CFD-PBPK track to construct a scatter plot and predict the fitting functions. The data reveals a logarithmic relationship between *I_o_*and *A*_ave_, proposing the equations “*I_o_ = 1.617In(A_ave_) + 7.703” and “I_o_ = 1.399In(A_ave_) + 7.396”* in case of normal breathing and sniffing profiles, respectively. Similarly, in case of adopting the time-averaged concentration, the curves could be shifted to “*I_o_ = 1.695In(C_ave_) – 5.910” and “I_o_ = 1.547In(C_ave_) - 3.894”.* The correlation coefficients indicate that the absorption flux and acetaldehyde concentration are reliable predictors of odor intensity, particularly during normal breathing. Additional investigations were conducted on the location-specific measurement of odor concentration to identify the appropriate spots that reflect the actual intensity of the odorants, either in the indoor environment, the breathing zone, or the inner regions of the human respiratory tract.

Fig 2A illustrates a detailed correlation analysis of odor intensity as a function of time-averaged equilibrium concentration at four distinct measuring spots: the centralized point in the fluid region of the yogurt headspace (Spot A), breathing zone (Spot B), nostrils (Spot C), and mucus layer in the olfactory wall (Spot D), as identified in Fig 2B. The predicted logarithmic fitting formulas are “*I_o_ = 1.166In(C_ave_) – 5.639”,* “*I_o_ = - 3.251In(C_ave_) + 24.066”,* “*I_o_ = 2.130In(C_ave_) - 13.875”, and* “*I_o_ = 1.695In(C_ave_) – 5.910”,* for the spots A, B, C, and D, respectively. The correlation coefficient R^2^ indicates that the olfactory region and nostrils capture the most reliable time-averaged concentration, reflecting perceived odor intensity. The rough sensory assessments of odorants in indoor environments or the breathing zone reflects negative/weak correlations. The dynamic change in the distribution of equilibrium concentration within the breathing zone (Spot B) showed an inadequate negative correlation with perceived odor intensity, potentially due to higher time-averaged concentrations during the exhalation phases, particularly in the first three cycles. Fig 2C further emphasizes the importance of location-specific measurements, presenting time-averaged concentration profiles across the measurement spots of five breathing cycles. A dynamic fluctuation of odor concentration in the nostrils and headspace was associated with dynamic changes in breathing airflow rate and the continued release of odor from the upper surface of the yogurt. However, relatively low and stable concentration profiles were observed in the breathing zone due to the strong turbulent flow that occurred at spot B, which maximized the dispersion of odorants to the outer regions. The mucus layer concentration profile indicated high sensitivity to low concentrations in the olfactory region (approximately 8% of the inhaled odor concentration).

**Fig 2.**
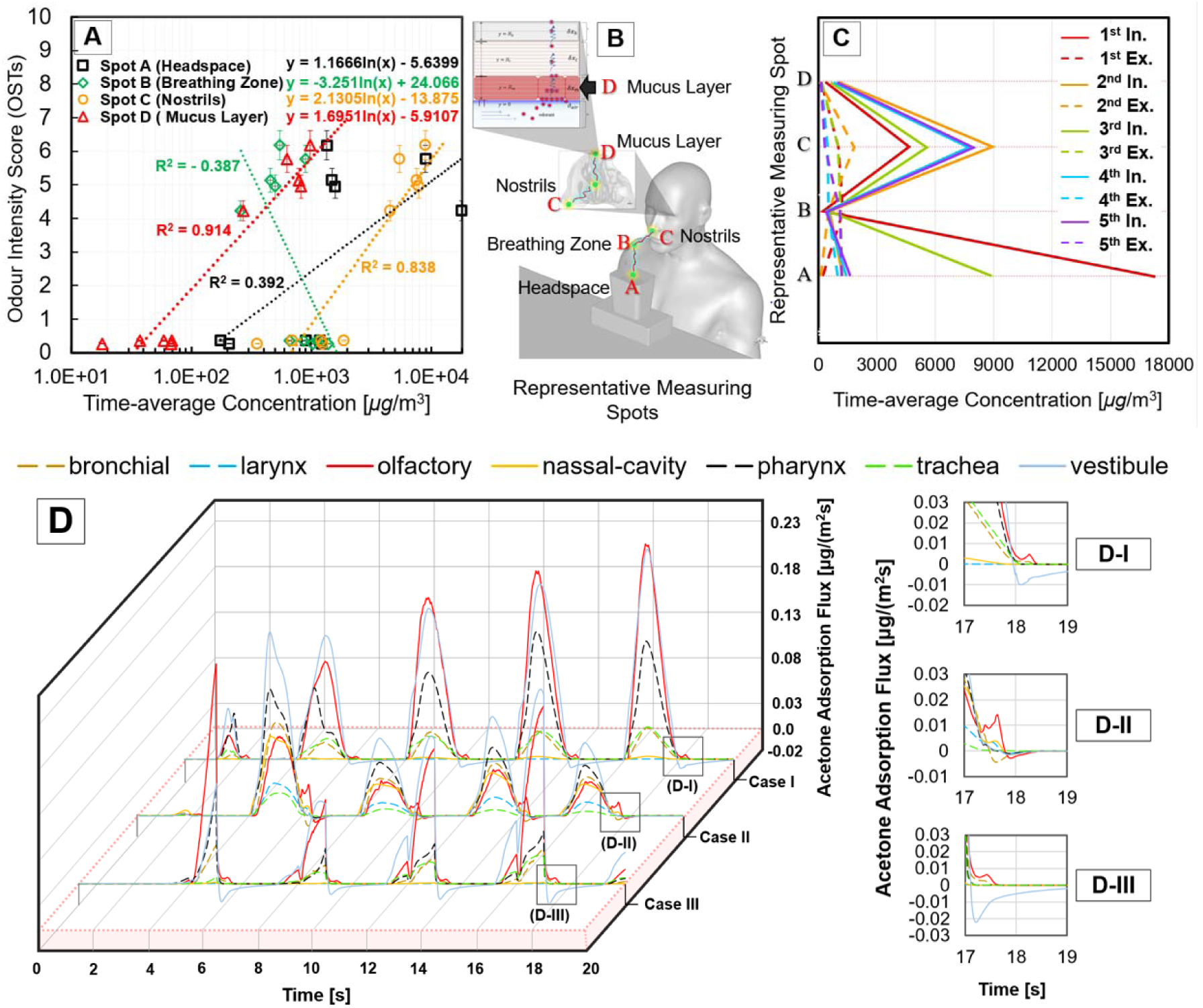
Observed olfaction phenomena and location-based analysis for accurate prediction of odor intensity/concentration. Panel (A) demonstrates the location-based analysis based on four main representative measuring spots as shown in panel (B), emphasizing the importance of using *in-silico* tools for accurate prediction of odor intensity as a function of chemical concentration in the appropriate spot. Panel (C) shows the dynamic change of acetaldehyde concentration profiles across different representative spots over five subsequent breathing cycles. Panel (D) clarifies the absorption flux trend of acetone, as one of the volatile organic compounds of the yogurt, into the mucus layer in three examined breathing profiles. The subpanels (D-I), (D-II), and (D-III) present a detailed view of the observed phenomena of absorption and desorption during the end of the inhalation phase and at the start of the subsequent exhalation phase, which represents case I, II, and III, respectively. The absorption flux was calculated on average for each airway segment.

Since acetone (C H O) is a benchmark VOC for assessing perceived odor intensity based on the ISO 16000-28 standard [30], we expanded the quantifications to include an analysis of advection-diffusion and intracellular kinetics of acetone within the respiratory wall (Fig 2D). In all three cases, the absorption flux peaks were predominantly observed in the olfactory region. Unlike acetaldehyde and other examined VOCs, acetone exhibited distinct desorption and absorption behaviors during the transitional timeframe. There was a sharp desorption of acetone immediately after the peak absorption, as indicated by the rapid decline in flux. This rapid desorption suggests a high turnover rate of acetone in the olfactory region, where the mucus surface releases acetone back into the air during exhalation (Figs 2D-I, 2D-II, and 2D-III). These phenomena are primarily attributed to the chemical properties of acetone, particularly its relatively lower partition coefficient between air and the mucus layer compared to other chemicals, coupled with its high diffusivity in both the air and mucus phases. However, owing to the low concentration of acetone in yogurt compared to acetaldehyde, these phenomena were observed at relatively minor values.

Visual observations through the CFD-PBPK track can enrich our discussion of the spatial transport of odorants within the breathing zone and inside the fluid airway domain (Fig 3). The simulation started with a constant concentration of VOCs in the headspace and upper-surface of the yogurt. Once the calculations were initialized, the odorants were immediately released from the headspace with a remarkable dispersion route directed toward the nostrils, which was induced during the first inhalation phase. At the end of this phase (2.0 seconds), the perceivable dose of odorants were dispersed omnidirectionally. At 3.0 seconds, the puff of the exhalation phase caused extensive scattering into a wider domain. Thus, the yogurt lost its condensed odorant concentration in the headspace far from the breathing zone, which was reflected in the low concentration trend of the first cycle, regardless of the breathing profile. During the second inhalation phase (5.0 seconds), the computer-simulated person (CSP) inhaled the diffused odorants near the nostrils and breathing zone, which occurred in the prior phase. This reflected the high concentrations present inside the nasal cavity and olfactory recess (Figs 3A, 3C, and 3E). During this time frame, the upper surface of the yogurt started to release the odor continuously, driven by the determined scalar value (Figs 3B, 3D, and 3F, and S6 Fig in the supporting information).

**Fig 3.**
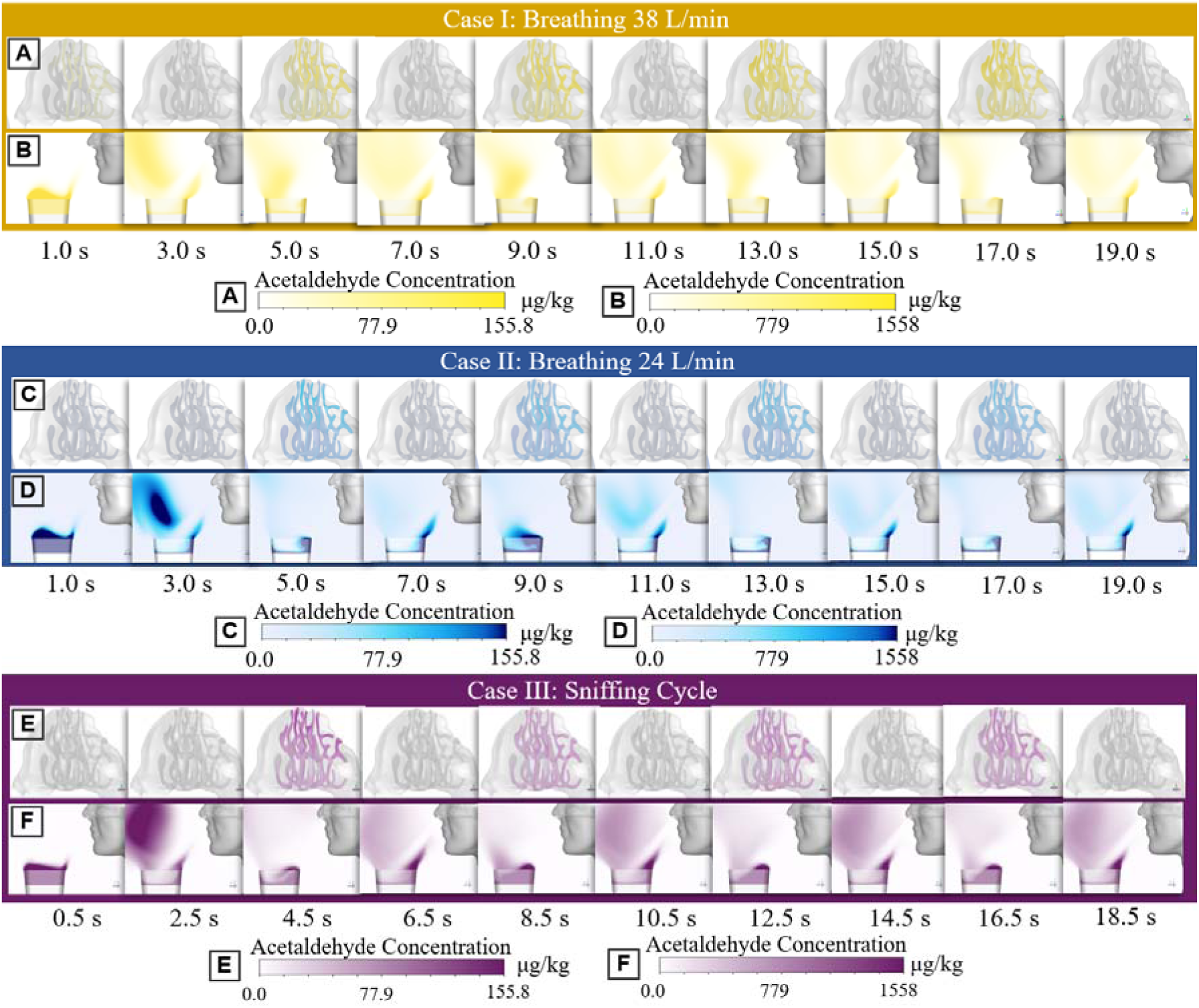
Visualizations of the acetaldehyde concentration distribution for three examined cases with different breathing profiles. Panels (A) and (B) show the concentrations inside the nasal passages and in the breathing zone for case I, respectively, at ten subsequent shots of the peaks of each inhalation and exhalation flow rate. The dispersion of the acetaldehyde from the headspace and surrounding the breathing zone can be easily identified through panel (B). Similarly, panels (C) and (D) represent case II, while panels (E) and (F) are for the sniffing cycle of case III, which has different peak times.

In conclusion, while OSTs provided average predictions of odor intensity through panelists’ perceptions, CFD-PBPK extended the quantification of absorption flux, approximated concentrations, and detailed the temporal and spatial transport of odorants into the olfactory recess in multiple and extended representative scenarios. Both integrated methods contribute to a comprehensive understanding of the olfactory process through odor sensory examinations. Exploiting the advantages of computational tools can maximize their potential in assessing olfaction and other respiratory phenomena using safe techniques. This approach offers potential for further computational and biomedical applications, such as developing sensitive electric noses, smart odor displays in virtual reality, smart food quality assessments, and other sensory implementations. Medically, it can enhance our understanding of the local fluid environment and the role of microbial ciliated tissues in the respiratory tract by investigating the microfluids.

## Materials and Methods

Two independent teams conducted odor sensory tests (OST) and CFD-PBPK simulations concurrently. The results remained confidential until both tracks were completed, culminating in the final analysis of odor sensitivity during the 20-second simulation period. S7 Fig illustrates the framework of the steps involved in each track, and highlights the connections established to ensure consistent boundary conditions. Key factors such as the ventilation flow rate, positioning of the yogurt and human subjects, air temperature control, and geometric characteristics were meticulously standardized across both tracks. The OST involved a rigorous training phase, whereas the CFD-PBPK track was validated and verified through mesh independence tests and other computational quality control methods, including the examination of various parameters. The integration of both approaches offers a novel pathway for accurate prediction of olfactory dynamics (S7 Fig).

### Setup of odor sensory tests (OST)

Initially, panelists attended comprehensive training sessions to calibrate their intensity rating and familiarize themselves with the OST procedures. Panelists were selected by excluding subjects with allergies or those who disliked yogurt. The experimental room was prepared to provide a stagnation region in the breathing zone to avoid fast dispersion of odorants (Figs 4A and 4B). Thus, the centralized area of the room was selected to conduct the OSTs with an average measured velocity of 0.02 m/s. The temperature and humidity were controlled at 23±2 °C and 50±5%, respectively. The gaseous odorants from the headspace of a Bulgarian plain yogurt sample were diluted in odor-sniffing bag samples that were used for the training stage (Fig 4C). The dilution rates of the gases were 3.16×, 10×, 31.6×, 100×, and 316×, labeled with numbers 10, 8, 6, 4, and 2, respectively, representing the strongest to weakest intensities. Thus, a 316× dilution rate was adopted as the odor-detection threshold. The panelists practiced odor intensity rating using nose pads attached to the sniffing bag samples and inhaled the gas to recognize each concentration of the samples until they felt proficient in evaluating the intensity (Fig 4C).

**Fig 4.**
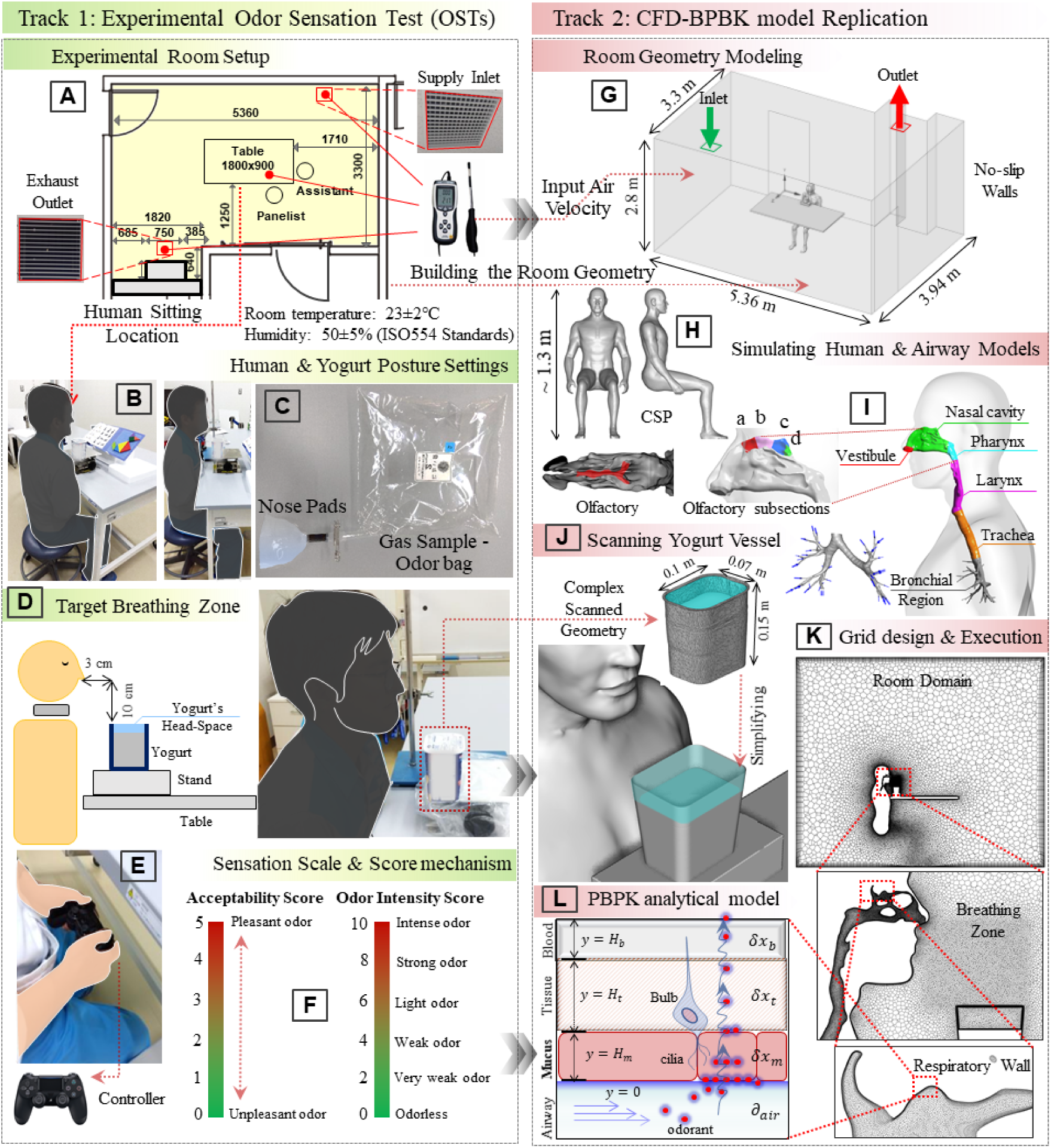
Methods and tools utilized in the odor sensory tests (OSTs) and computational fluid dynamics and a physiologically based pharmacokinetic (CFD-PBPK) model. Panels (A), (B), (C), (D), (E), and (F) illustrate the dimensions of the experimental room, the posture setting of the human and yogurt vessel, the gas sample bags and nose pads used in the training phase, the identification of target area, the controller, and lastly, the scale of odor intensity and acceptability as a benchmark for scoring, respectively. In the CFD-PBPK tract, panels (G), (H), (I), (J), (K), and (L) present the room geometry, the seated computer-simulated person (CSP), the segments of the upper airway mode, the yogurt vessel, the grid design in different locations, and lastly, the schematic diagram of the PBPK model integrated into the respiratory wall, respectively.

The yogurt vessel was fixed close to the human face, 10 cm below the nostrils, and at a horizontal distance of 3 cm (Fig 4D). During the OSTs, the score input device was a DUALSHOCK 4 (Sony Interactive Entertainment Inc., Sony, Japan) connected via Bluetooth to a Windows PC for scoring at a fast frequency (Fig 4E). Two respiratory profiles were tested: normal breathing with two-second inhalation and two-second exhalation, and a sniffing profile with one-second inhalation and three-second exhalation, respectively. The panelists rated the intensity and acceptability of five breathing cycles (20 seconds) based on the benchmark scales (Fig 4F). Each panelist evaluated eight OSTs, consisting of a set of two evaluations with two different sniffing patterns, and repeated these four OSTs twice.

The experimental room (27 m^2^) had a mixed ventilation mode with a small supply inlet and exhaust outlet allocated to the opposite corners of the ceiling. However, to prevent exposure of direct airflow, the panelists were seated in the center of the room. A table was used to fix the position of the yogurt vessel close to the human and block the accelerated airflow from the ground to the upper region. These setup conditions ensured an air stagnation region in the breathing zone to avoid quick dispersion of odorants. Simultaneously, the geometry of the CFD track was built based on the physical and airflow measurements of the experimental room. The room inlet velocity was kept at 0.51 m/s, the supply temperature at 23 °C, and turbulence intensity maintained at 10%, respectively. The additional boundary conditions are summarized in S2 Table.

#### Discretization

For the model discretization, the suitable grid design for the computational load at the cost of accuracy contains 9.5 million cells, of which 3.6 million cells were located in the airway fluid regions, and 2.3 M in the breathing zone and headspace of the yogurt vessel (Fig.4K). A multi-layer model of air-mucus-tissue-blood (AMTB) proposed by Tian and Longest (25) was employed in the airway wall to analytically express the absorption and concentration of yogurt VOCs (Fig 4L). Extended clarifications of the materials and methods are provided in the SI Appendix. A grid independence test and velocity profile checkup was conducted to ensure the applicability of the model for approximating the fluid regions, according to previously reported validations [31,32,33,37]. The tested grid designs were illustrated in Fig. S8. A refined prism boundary layer was created with a thickness of less than 0.1 mm to approximate near-wall y+ values of less than 1 at the maximum inhalation/exhalation airflow velocities. All the governing equations for continuity, energy, and scalar transport (odorant dispersion) were resolved using the shear stress turbulence model (SST k-omega). Various engineering applications have confirmed the effectiveness of these methods in terms of cost and time efficiency in similar engineering applications [15,38,39]. This model can resolve the velocity profile in the viscous sub-layer of the interior airway wall. For the convection term, the second-order upwind scheme was selected for all governing equations except for pressure, and the standard scheme was preferred.

#### Examined breathing profiles

In the numerical model of the respiratory tract, the airflow distribution across 38 upper airway compartments was estimated using the structural models proposed by Shelley et al. [40] and Horsfield et al. [41]. The inlet and outlet velocities for each compartment were predicted based on the cross-sectional diameters of the channels and their positions within the airway tree. The temperature of the internal airway surfaces was maintained at a constant 34 °C. For the CSP wall, the skin temperature was calculated following a fixed routine, aligned with the model proposed by Fanger [42,43], using a heat balance equation between the core and surrounding temperatures. This method demonstrated stable predictions during the simulation training set [29,44]. S9 Fig presents a comparative analysis of the three different breathing profiles, where variations in the inhalation and exhalation flow rates were evident in the velocity distributions (S9-A and S9-B). Correspondingly, the dispersion zones of the yogurt odorants were distinguishable as well (S9-C).

#### Extended description of the PBPK model

A multi-layer air-mucus-tissue-blood (AMTB) model proposed by Tian and Longest [25,45,46] was employed to analytically express the absorption and concentration of yogurt VOCs. Odorants deposited from the airway gas phase to the airway wall are transported by diffusion within the mucus layer (H_m_=15 μm), the mucosal epithelial layer (tissue) (H_t_=50 μm) and the sub-epithelial blood layer (H_b_=15 μm). The surface concentration was calculated at the air-mucus boundary using the absorption equilibrium expressed in Eq. (1) and the diffusion flux conservation equation for the low-viscosity layer in Eq. (2). Thus, the mucus wall surfaces were assumed to be no-slip walls.

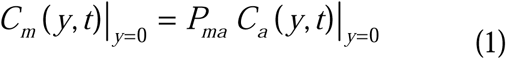

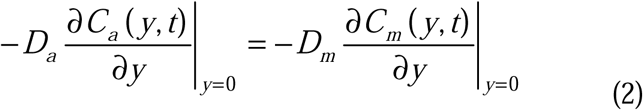

where *C_a_ and C_m_* refer to the chemical concentration in the gas phase and mucus layer [μg/m^3^], respectively. *D_a_* and *D_m_* are the diffusion coefficients of each chemical in the gas phase and mucus layer [m^2^/s], respectively. *P_ma_* is the partition coefficient [-] between the concentration in the air layer and the concentration in the mucus layer. All these parameters are summarized for each VOC in S3 Table.

Accordingly, the absorption equilibrium and flux conservation between the mucus layer and mucosal epithelial (tissue) layer interface are expressed in Eq. (3) and (4), respectively:

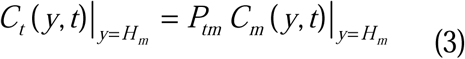

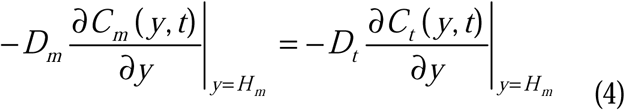

where *C_t_* refers to the chemical concentration in tissue layer [μg/m^3^]. *D_t_* is the diffusion coefficient of each chemical in the same layer [m^2^/s]. *P_tm_*is the partition coefficient [-] between the concentration in mucus and tissue layers.

Similar to the air-mucus boundary, we assumed local equilibrium at the mucus-mucosal epithelium and mucosal epithelium-subepithelial blood boundaries with an equal concentration of zero for each layer in the initial conditions, as expressed by Eq. (5)

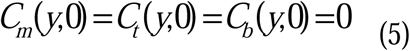

In conjunction with the airflow field computation, the scalar transport equation was solved using Eq. (6)

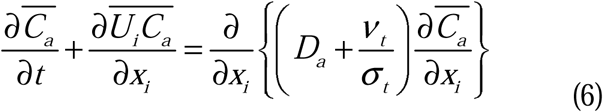

where *C_a_* is the concentration of the chemicals in the gas phase [μg/m^3^], ensemble averaged for the Unsteady Reynolds Averaged Navier-Stokes Equation (URANS) model analysis. *D_a_* is the molecular diffusion coefficient of the chemical in the air [m /s], *ν_t_* is the eddy kinematic viscosity coefficient calculated by the RANS model, and *σ_t_* is the turbulent Schmidt number.

#### Approximation of the chemical physical properties in CFD-PBPK model

The partition coefficients between the air and mucus layers (*P_a:m_*) were calculated using Henry’s law of solubility constants (*H*), as extensively reported for a wide range of chemicals by Sander [47]. Since the reported values were at 25 °C, they were converted to the equivalent value at 36 °C (represents the respiratory wall temperature using the Van’t Hoff equation for temperature dependence of equilibrium constants. The diffusion coefficients of chemicals in mucus and partition coefficients at the air-mucus boundary are provided for mucus composed primarily of water (95-99%), a mixture of proteins, mucins (a type of glycoprotein), DNA, cellular debris, etc. [18,23,48]. This correction requires knowledge of the octanol-water partition coefficients of each chemical, which were obtained using the KOWWIN model at the EPI Site [18]. Therefore, the values of the diffusion coefficient in water and the Partition Coefficient at the gas-water boundary were adopted for approximation. The diffusion coefficients of the chemicals in the tissue epithelium were assumed to be proportional to one-third of the diffusion coefficient in the mucus layer. The diffusion coefficients of the chemicals in the subepithelial layer through which the blood flowed were calculated using the Stokes-Einstein rule. The viscosity coefficient of blood was set to 4.0×10^-3^ Pa second.

### Properties of yogurt volatile organic compounds (VOCs)

The sources of odorants in daily life are diverse. Some of them can be harmful even with brief or prolonged exposure. Panelists were asked to notify us of irregular physical conditions on the test day with obligatory one-hour fasting before execution to maximize their sensitivity to volatiles. Food safety investigations on yogurt odorants have reported the appropriate safe range for daily consumption. In previous investigations of the key odor-active compounds in yogurt, 13 main odorants were detected, of which all the examined odorants in the CFD-PBPK track were involved [49]. Acetaldehyde is a highly water-soluble chemical, toxic to airway tissue epithelia [24]. Unlike multiple pollutant sources of acetaldehyde, yogurt contains a relatively minimal concentration (average between 10 and 50 ppm, depending on the product) that is safe for short-term exposure. However, the frequent intake of such chemicals from multiple sources may pose a potential risk. Moreover, based on various reported studies, 2,3-butanedione and 2-heptanone in yogurt are the most vital components with the highest flavor dilution factor [49,50,51]. In the current study, the yogurt used in the OST consists of 10.6 ppm acetaldehyde (400 g vessel), which was the dominant chemical. The measurements showed wide range of chemical concentrations and the most common chemicals for evaluation in CFD application were selected. These were acetaldehyde (CH_3_CHO), acetone (C_3_H_6_O), 2,3-butanedione (C_4_H_6_O_2_), 2,3-butanediol (C_4_H_10_O_2_), acetic acid (C_2_H_4_O_2_), and 2-heptanone (CH_3_(CH_2_)_4_COCH_3_). The concentrations and dispersion rates of the chemicals in the yogurt vessel was measured to reflect the actual boundary conditions of the CFD simulations. In the dead-space and the upper-surface of the yogurt, the average chemical concentrations were set at 10.6, 0.35, 0.18, 0.06, 0.05, and 0.01 ppm for acetaldehyde, acetone, 2,3-butanedione, 2,3-butanediol, acetic acid, and 2-heptanone, respectively. S3 Table summarizes the properties of each odorant from the different sources. The diffusion coefficient in the air (*D_a_*) where either cited from experimental data (if available) or estimated based on the online US-EPA tool for Site Assessment Calculation. In OST sessions, various yogurts with different storage conditions were used to accommodate a wide range of yogurt evaluations. Test samples included yogurts used within eight days of production.

## Acknowledgments

This study was approved by the Ethics Committee of the Meiji Institutional Review Board (no. 230, approved on October 12, 2023, UMIN: UMIN000052536).

## Supporting Information

**S1 Fig. Average concentration distribution of the yogurt’s VOCs in the mucus layer during 20 seconds of computational fluid dynamics (CFD) simulations.** Quantifying local-based analysis of four subsections of the olfactory wall (namely, olfactory “a,” “b,” “c,” and “d,” from the most anterior to the most posterior sides of the olfactory recess, respectively). (A) shows the results of case I, (B) case II, and (C) case III. The dominance of acetaldehyde as a key active component of yogurt is evident.

**S2 Fig. Average concentration distribution of the yogurt’s sub-active VOCs (2,3-butanedione, 2,3-butanediol, acetic acid, and 2-heptanone) in the mucus layer during 20 seconds of CFD simulations.** Results from case I (A & B), case II (C & D), and case III (E & F). Due to the special properties of 2-heptanone, which has high molecular weight (MW) and low diffusion coefficient in different phases, it showed irregular concentration distribution, unlike other chemicals.

**S3 Fig. Absorption flux of the yogurt VOCs into the mucus layer over seven segments of human upper airway walls in the breathing profile case I.** The results represent (A) acetaldehyde, (B) acetone, (C) 2,3-butanediol, (D) 2,3-butanedione, (E) acetic acid, and (F) 2-heptanone.

**S4 Fig. Absorption flux of the yogurt VOCs into the mucus layer over seven segments of human upper airway walls in the breathing profile case II.** The results represent (A) acetaldehyde, (B) acetone, (C) 2,3-butanediol, (D) 2,3-butanedione, (E) acetic acid, and (F) 2-heptanone.

**S5 Fig. Absorption flux of the yogurt VOCs into the mucus layer over seven segments of human upper airway walls in the sniffing profile case III.** The results represent (A) acetaldehyde (B) acetone, (C) 2,3-butanediol, (D) 2,3-butanedione, (E) acetic acid, and (F) 2-heptanone.

**S6 Fig. Temporal and spatial distribution of yogurt’s volatile organic compounds (VOCs) as outputs of CFD-PBPK track in the examined case II.** (A) Comparison of the dispersion patterns between six different VOCs, including acetaldehyde (CH_3_CHO), acetone (C_3_H_6_O), 2,3-butanedione (C_4_H_6_O_2_), 2,3-butanediol (C_4_H_10_O_2_), acetic acid (C_2_H_4_O_2_), and 2-heptanone (CH_3_(CH_2_)_4_COCH_3_). The diffusion coefficient in air is a fundamental parameter for assessing the dispersion rate of the chemicals in the breathing zone and releasing patterns from the upper-surface of the yogurt into the headspace, directed to the nostrils. (B) Timeline and detail during the first exhalation phase, from 2.05 seconds to 3.75 seconds. It reveals the fast dispersal of the condensed odorants in the headspace, induced by the high flow rate of expiratory jet.

**S7 Fig. Framework of the research methodology.** Divided into two sections, track 1 represents the odor sensory tests (OSTs), and track 2 represents the computational fluid dynamics/physiologically based Pharmacokinetic Model (CFD-PBPK). Both tracks were involved in the broad understanding of the temporal and spatial olfaction dynamics through the integrated outputs. Each step was conducted carefully, through preparation, replication, and validation to ensure accurate predictions. All correlations between the tracks were illustrated.

**S8 Fig. Tested grid designs that were used to ensure mesh independence.** The figures show four constructed models from A to D, referring to the coarse mesh to the highly refined mesh designs, respectively.

**S9 Fig. Visualization of the captured CFD-PBPK spatial concentration distribution.** (A) Airflow velocity counter distribution in the breathing zone during the peaks of the respiratory and expiratory phases in different breathing profiles. (B) Examined transient breathing profiles. (C) Three dimensional illustrations of odor dispersion zones, assuming iso-surface region of the acetaldehyde concentration higher than 15.5 μg/kg, representing equal or less than 1% of the initial concentration in the headspace of the yogurt vessel.

**S1 Table. Output averaged data from the odor sensory tests.** The data summarize the average perceived odor intensity by 11 panelists.

**S2 Table. Boundary conditions and solving models of the CFD-PBPK track.** The table summarizes the settings for preparation of the inlet, outlet, and walls boundary conditions for different objects of the CFD model.

**S3 Table. Properties of yogurt VOCs utilized in developing the PBPK model.** The data include the partition and diffusion coefficients for the examined six gas-phase chemicals in the air, mucus, tissue, and blood layers.

